# A Rational Drug Combination Design to Inhibit Epithelial-Mesenchymal Transition in a Three-Dimensional Microenvironment

**DOI:** 10.1101/148767

**Authors:** Farnaz Barneh, Mehdi Mirzaie, Payman Nickchi, Tuan Zea Tan, Jean Paul Thiery, Mehran Piran, Mona Salimi, Fatemeh Goshadrou, Amir R. Aref, Mohieddin Jafari

## Abstract

Epithelial-Mesenchymal Transition (EMT) is a major player of tumor invasiveness whose inhibition is challenged by redundancy of multiple inducing factors. We applied a systems-pharmacology approach by integrating network-based analyses with multiple bioinformatic resources to design a drug combination regimen reversing EMT phenotype in aggressive cancers. We observed that histone deacetylases were critical targets to tune expression of multiple epithelial versus mesenchymal genes. Moreover, SRC and IKBK were the principal intracellular kinases regulating multiple signaling pathways. To validate the anti-EMT efficacy of the target combinations, we inhibited the pinpointed proteins with already prescribed drugs and observed that whereas low dose mono-therapy failed to limit cell dispersion from collagen spheroids in a microfluidic device as a metric of EMT, the combination fully inhibited dissociation and invasion of cancer cells toward co-cultured endothelial cells. Given the approval status and safety profiles of the suggested drugs, the proposed combination set can be considered in clinical trials.

## 1. Introduction

Epithelial-Mesenchymal Transition (EMT) is a complex reprogramming process through which carcinoma cells facilitate their dissemination, acquire stemness features and drug resistance [1, 2]. Despite recognition of EMT as the key machinery to evade environmental limitations in tumors, full understanding of the process is yet far from complete and accordingly very few clinical trials used EMT as a biomarker [3-5]. Developing therapeutic approaches to inhibit this phenotype is also hampered by the dynamic nature of EMT necessitating a new rational design to hit multiple underpinning targets but the challenge then would be to select drug combinations that would be likely to show efficacy in patients [6].

Many targeted therapies, primarily developed to inhibit component of activated pathways in cancer are currently being tested for their anti-EMT effects. For instance, SRC and MEK inhibitors, small molecules and antibody inhibitors of growth factor receptors have shown promising anti-EMT effects [7, 8]. However, from a clinical perspective, their efficacy is hindered by the development of resistance making it an urgent need to combine the existing disparate and yet conflicting knowledge of EMT systematically to expedite the translational steps of selecting efficient drug combinations.

To tackle this complexity, high-throughput experiments have begun to shed light on details of various stimuli and mechanisms leading to EMT [9-16]. We are thus witnessing an increasing number of published gene lists demonstrating EMT portrait in aggressive cancers. Efforts have been also made in performing meta-analyses to integrate a number of expression datasets with several EMT inducers to provide a more unified description of EMT irrespective of nuance differences in the experimental conditions [17, 18]. These signatures however, share insignificant direct gene overlap and such variations complicate selection of appropriate therapeutic targets.

We believe that a rational integration of numerous bioinformatic and statistical tools are required to address the aforementioned problems, so that the biological researchers would be able to perceive principal mechanisms of EMT as a whole [19] and [20]. Herein, we aim to set a stage for assimilation of multiple bioinformatic resources to propose a therapeutic multi-target shotgun against EMT from currently published signatures (Figure 1). With commitment to simplicity, we performed sequential systems-based in silico experiments to hypothesize the essential targets that if hit together by co-drugging, will inhibit hallmarks of EMT in aggressive cancers. The anti-EMT efficacy of the proposed drug combination showed to be promising in a 3-dimensional (3D) co-culture model of EMT recapitulating cancer cell interactions with microenvironment. It is noteworthy to mention that while the key findings of our straightforward approach were in line with previous literature derived from years of conventional wet lab research; such approach is of paramount importance in transfiguring segmented data into a hypothesis-inspiring pipeline energizing another round of improved drug combination design for clinical research.

**Fig.1.**
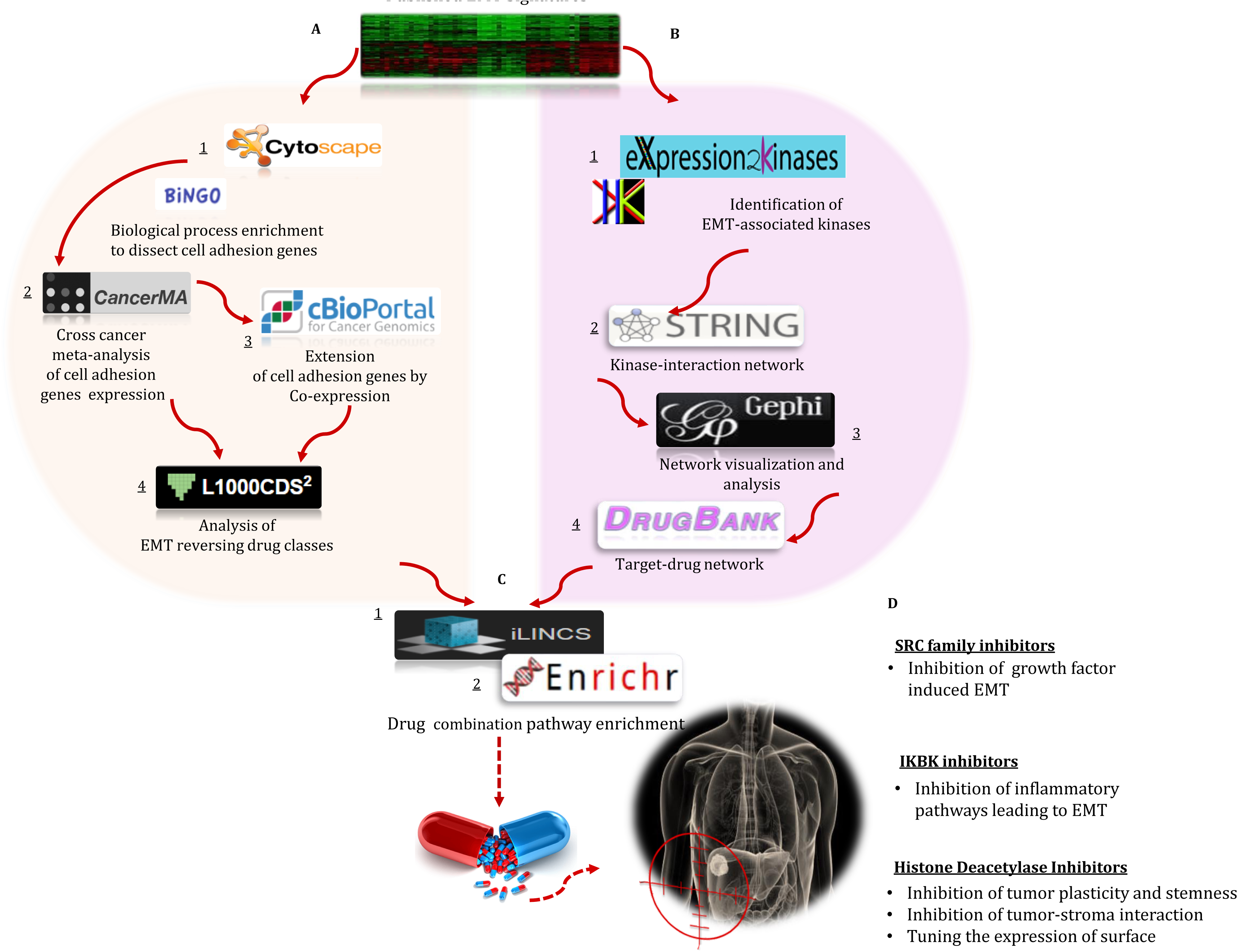
A systems-pharmacology workflow with integration of multiple bioinformatic tools sheds lights on key players of EMT in cancer. **(A)** From published EMT signatures, genes associated with cell adhesion were extracted (BiNGO App in Cytoscape), converted to an up/down signature in various solid cancers (CancerMA) and extended by co-expression (cBioPortal) to identify drugs (L1000CDS^2^) that induce expression of epithelial-related genes while downregulate mesenchymal associated genes. **(B)** EMT-associated kinome were identified by enrichment analysis (Expression2Kinases) and were used to create a kinase-Interaction Network in EMT (STRING). Network analysis (Gephi) was performed to identify important kinases and their inhibitors (DRUGBANK). **(C)** The signatures of the identified drugs were extracted (iLINCS) and subjected to pathway enrichment analysis (Enrichr) to estimate drug combination efficacy. **(D)** Summary of drug classes in combination and their prospecting effects which is hypothesized to inhibit multiple aspects in EMT.

## 2. Materials and methods

### 2.1. Compilation of published EMT signatures

EMT-associated gene signatures were obtained from the literature sources including Byers et al, [21] and Tan et al. [13]; also from two meta-analyses performed in 2012 [17] and 2016 [18]. The gene lists in the four aforementioned signatures were integrated to create an EMT seed gene library.

2.2. Enrichment-based and analytical tools

Enrichment tests were performed in ***“BiNGO”*** Cytoscape plugin [22], Gene set enrichment analysis (GSEA) in ***“Molecular Signatures Database (MSigDB) version 6.0”*** (software.broadinstitute.org/gsea/msigdb) [23] and also Enrichr (amp.pharm.mssm.edu/Enrichr) web platform as of Feb 2017 [24]. ***“CancerMA”*** database (www.cancerma.org.uk/information.html) accessed on Feb 2017 was used to perform meta-analysis across 80 cancer datasets for 13 different cancer types [25]. ***“L1000CDS^2^”*** search engine (amp.pharm.mssm.edu/L1000CDS2) accessed date on Feb 2017 was used to query L1000 datasets and identify EMT reversing drugs [26, 27]. ***“Expression2Kinases (X2K)”*** downloaded in Nov 2016 (www.maayanlab.net/X2K) was used to identify EMT-associated kinome which utilizes enrichment analysis of kinases and their substrates [28]. Initially, a bottom-up approach was taken starting from identification of transcription factors as described previously in [29] and [30] which mainly identified intracellular non-receptor kinases linking signal transduction pathways to gene expression changes in the nucleus. Individual EMT gene signatures were also directly submitted to KEA module of X2K to identify kinases regulating protein functions at the extracellular and cell membrane levels. ***“DRUGSURV”*** database accessed on March 2017 (bioprofiling.de/GEO/DRUGSURV) was used to reveal the effect of drug targets on survival [31]. ***“CellMiner”*** analysis tool version 2.1 [32] (discover.nci.nih.gov/cellminer) and ***“CSIOVDB”*** database of ovarian cancer microarray gene expression version 1.0 [33] (csibio.nus.edu.sg/CSIOVDB/) were applied to query drug targets expression analysis across NCI-60 cancer cell line panel and ovarian cancer datasets.

### 2.3. Databases, public resources and software

***“cBioportal”*** (www.cbioportal.org) database version 1.X.0 was used to expand differentially expressed genes in each cancer with The Cancer Genome Atlas (TCGA) data sets by performing co-expression [34]. Random gene lists were obtained x201C;Random Gene Set Generator"embedded.in.molbiotools.(http://www.molbiotools.com/randomgenesetgenerator.html). ***“STRING 10.0”*** (https://string-db.org) database [35] was used to convert the kinase lists into a kinase-kinase interaction network. Analysis of proper network centrality metrics was performed and visualized in ***“Gephi 0.8.0”*** [36]. ***“CREEDs”*** database version 1.0 (amp.pharm.mssm.edu/CREEDS/) was queried and GEO IDs including GSE37428, GSE15161, GSE17511, GSE26410 were extracted which represented SRC and IKBK overexpression [37]. To access drug-related signatures, the ***“iLINCS”*** web platform (http://ilincs.org) was queried to extract datasets of Vorinostat (LINCSCP_34467 and LINCSCP_66500), Dasatinib (LINCSCP_36498 and LINCSCP_6177) and Sulfasalazine (LINCSCP_34467 and LINCSCP_5717) in MCF-7 and A549 cell lines. Expression of drug signatures in murine hematopoietic cells was analyzed with “my gene set” tool in “ImmGen” project (immgen.org) [38].

### 2.4. Experimental validation of anti-EMT effects of predicted drugs

All procedures to co-culture A549 spheroids with HUVECs in 3-dimensional (3D) microfluid devices were performed as previously described in [39, 40] Drugs chosen based on in silico predictions, were dissolved in DMSO with final concentration of 1µM with media and injected into the channel inlets either alone or in combination. Concentration of drugs was selected based on their relevance to achievable clinical doses. High content imaging was performed after 0, 12, 36, 60 and 84 hrs via FluoView 1000 confocal microscopy (Olympus, Japan). The extent of cell dispersion from the spheroids was quantified by measuring mean distance between red pixels and was normalized by dividing to the mean distance in zero time point.

### 2.5. Statistical analysis

For enrichment tests FDR-adjusted P-value <5%, for meta-analysis and -1.5 <fold change <+ 1.5 and in co-expression analyses, Pearson’s correlation co-efficient >0.8 were considered significant. In cell-based experiments two tailed t-test with statistical significance of was level < 0.05 in GraphPad Prism 6 (GraphPad Software; La Jolla, CA). Quantification of cell-based experiments was performed with MATLAB 2014b (MathWorks; Natick, MA).

## 3. Results

### 3.1. Unified EMT gene set library expression in cancers

EMT-related signatures were compiled from four published studies (for details see methods) and a library of 962 genes was generated. Taking into account the importance of cell adhesion-related genes in initiation of EMT, the biological process gene cluster under “Cell-adhesion, GO:0007155” term was dissected from the main library resulting in a subset of 93 genes. The status of their differential expression was assessed by performing meta-analysis across eighty datasets and thirteen cancer types using ***“CancerMA”*** database. Complete list of differentially regulated cell-adhesion genes across multiple cancers is provided in (**Supplementary File 1**). This approach also allowed converting the gene set into an upregulated (regarded as mesenchymal-related genes) and downregulated (regarded as epithelial-related genes) signature in a cancer-dependent manner. Eighty two genes were found to be differentially deregulated in multiple solid cancers however, no common cell-adhesion related gene was found to be deregulated in all cancer types of the study; e.g. a metalloproteinase (ADAM12) and Fibronectin (FN1) were upregulated only in five cancer types (**Figure 2A**) while FBN1 Fibrillin (FBN1) was downregulated in seven cancer types and Dermatopontin (DPT) and stromal cell-derived factor-1 (CXCL12) were downregulated in six and five cancer types respectively (**Figure 2B**).

**Fig.2.**
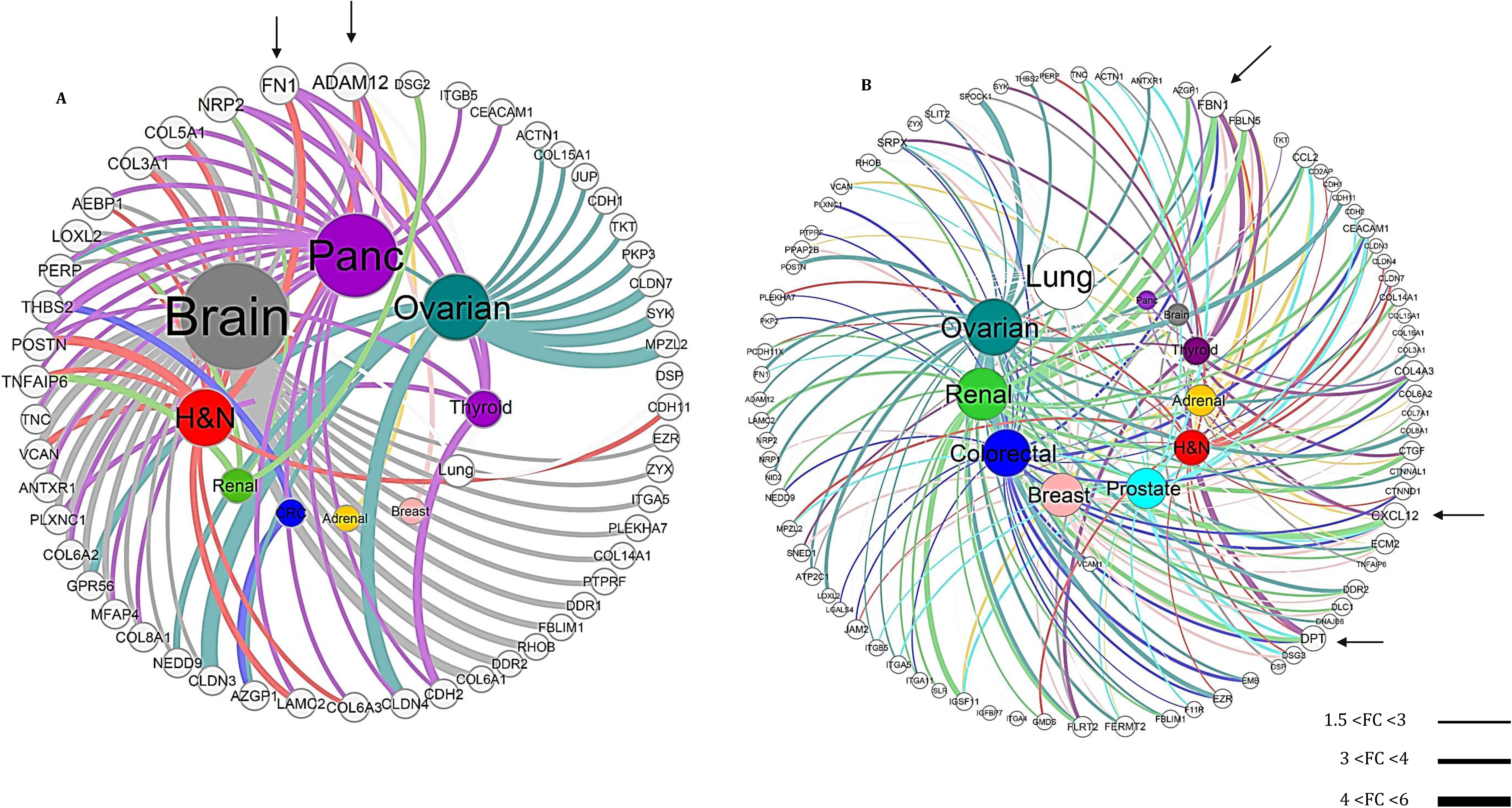
Meta-analysis shows inconsistent expression of cell adhesion-related genes across various solid cancers. **(A)** Upregulated cell adhesion-related genes dissected from collated EMT signatures. Expression status was determined by performing meta-analysis in CancerMA database for different cancer types. These upregulated genes were considered as mesenchymal genes **(B)** Downregulated cell-adhesion related genes from collated EMT signatures in various cancers obtained by meta-analysis. These downregulated genes were assumed to be associated with epithelial characteristics. The outer layer shows de-regulated genes and the inner layer represents cancer types. Edge thickness in the networks signifies folds change (FC) range. ±1.5 FC was considered significant. Arrows indicate most similar genes in different cancers.

### 3.2. Prediction of drugs to tune expression of cell adhesion-related genes

To identify drugs that reverse the signature of cell-adhesion related genes in multiple cancers, the up/down genes for each cancer were submitted into ***“L1000CDS^2^”*** search engine to prioritize signature reversing drugs based on the overlap score calculated by characteristic direction method [41]. We observed that even though each cancer exhibited different set of expression patterns for cell-adhesion related genes, Histone Deacetylase Inhibitors (HDACIs) including Trichostatin A, Vorinostat, Panobinostat and Pracinostat were repeatedly detected to reverse the expression of various genes across ten cancers in the study **(Figure 3A)**. Since HDACIs induce significant changes in gene expression, a P-value from z-test was computed to compare the overlap score of HDACs signatures with that of fifty random gene lists with the same length of up/down genes for each cancer type to ensure their overrepresentation in the results was due to their effect on reversing EMT-associated genes more significant than a random set. Except for adrenal cancer, HDACIs significantly reversed EMT-associated gene sets in ten solid cancers **(Figure 3B)**. Intermediate states of EMT are associated with deregulation of varying degrees of epithelial and mesenchymal genes in the cell surface. To further extend these deregulated genes with patient data across multiple cancers, the selected upregulated genes for each cancer were used as seed and were expanded with Cancer Genome Atlas (TCGA) datasets (**Supplementary File 2**) using co-expression (Pearson’s r>0.8) toolbox in <***“cBioPortal”*** web tool [34]. In resubmitting the expanded gene lists to L1000CDS2, HDACIs were observed among the top ranked drugs to reverse expression pattern of the extended signatures in eight cancers which are marked with asterisk in **Table 1**

**Fig.3.**
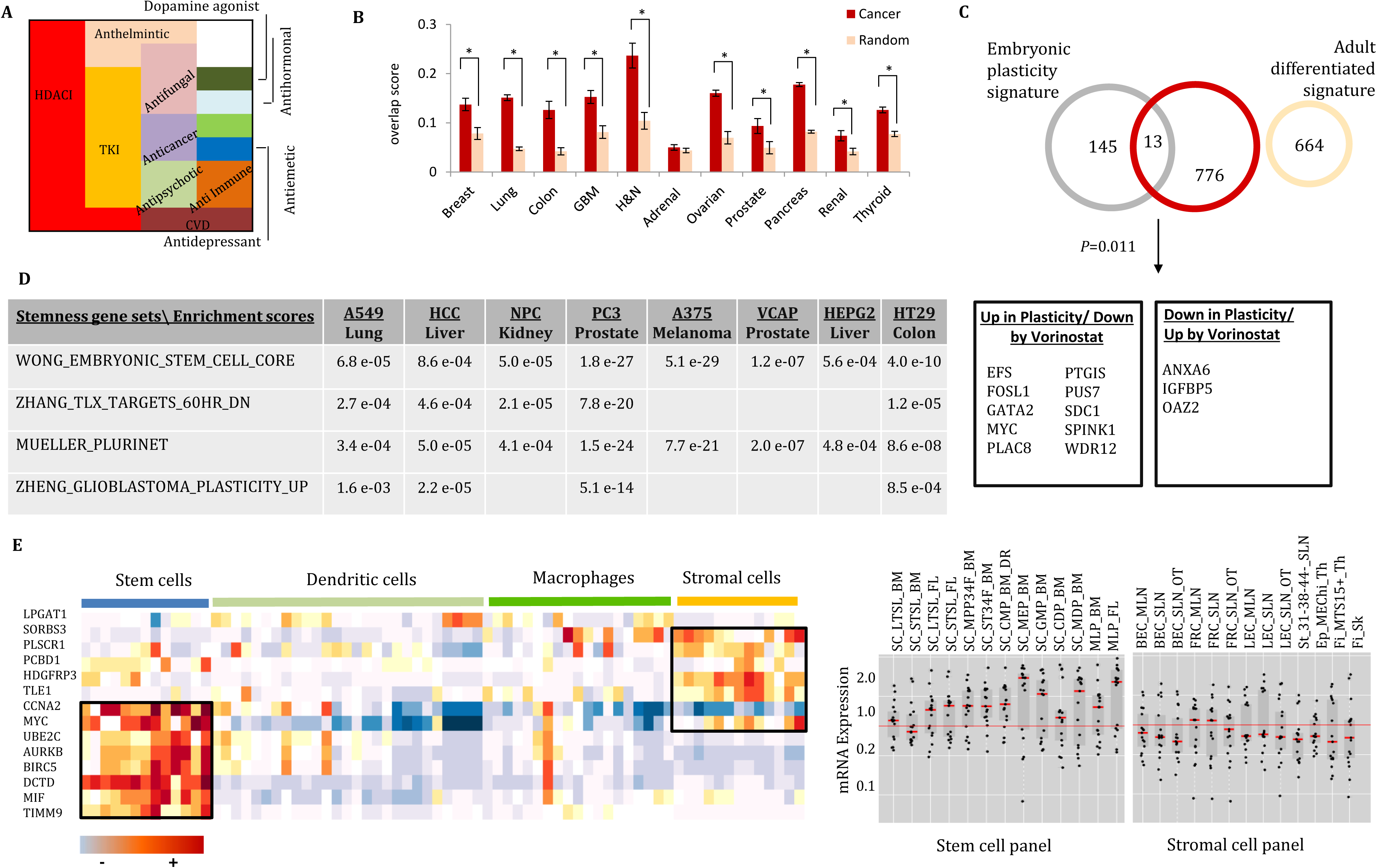
HDACIs are a notable class of drugs to reverse multiple EMT-associated processes. **(A)** Relative illustration for number of cancer types whose EMT induced cell-adhesion related genes sets are reversed by each class of drugs. **(B)** Efficacy of HDACIs to reverse EMT-associated gene signatures in various cancers based on characteristics direction signature score compared to random gene sets with the same length (z-test, * p<0.05). **(C)** Overlap between up/down regulated genes in embryonic plasticity and adult tissues which are reversed by Vorinostat. **(D)** Gene set enrichment analysis of differentially expressed genes induced by Vorinostat in multiple cell lines with stem cell signatures in MsigDB database version 6.0. (E) Heatmap and box-plot mRNA expression of significantly downregulated genes by vorinostat in various mouse hematopoietic and stem cell lines in ImmGen project.

**Table-1.**
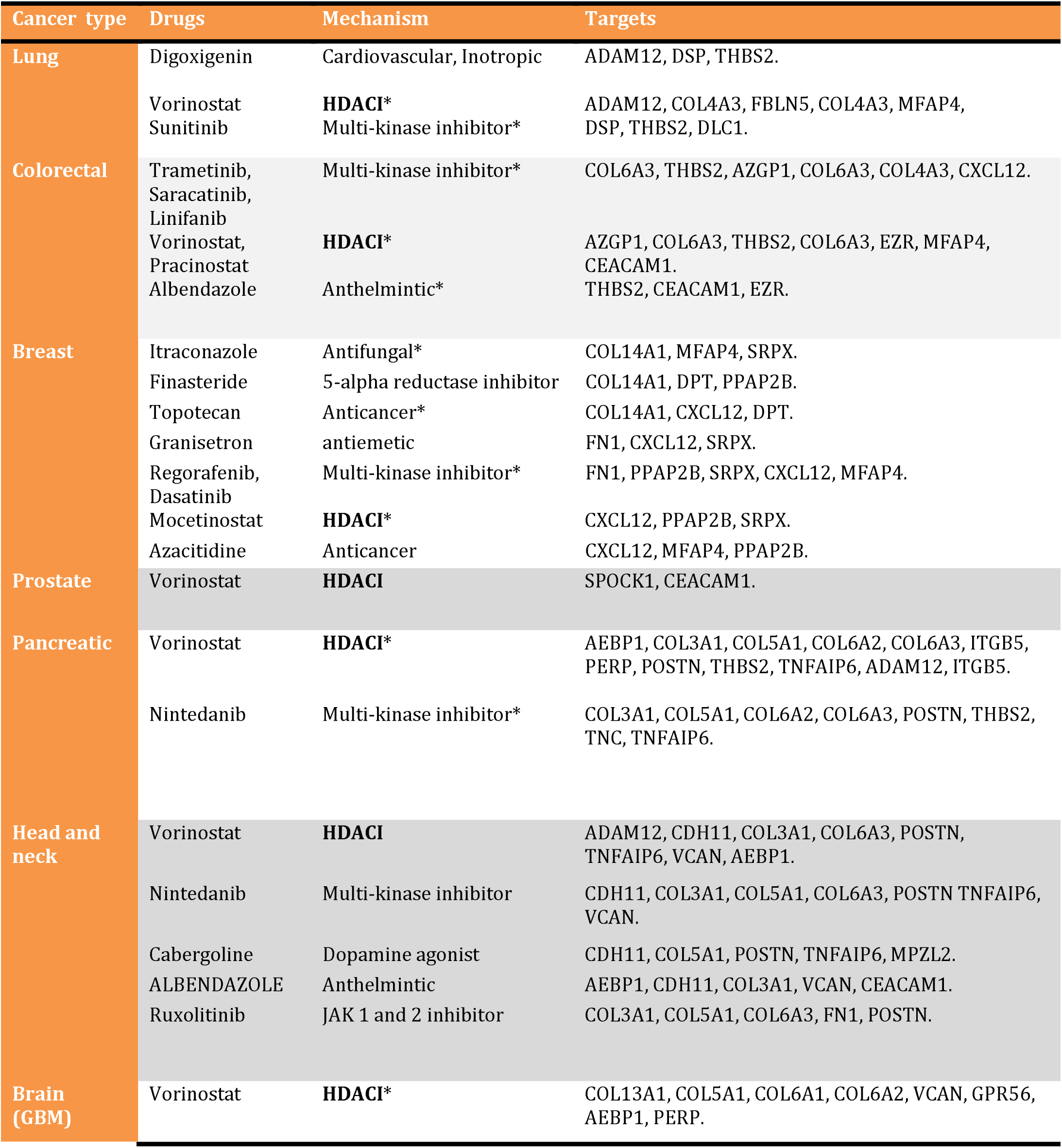
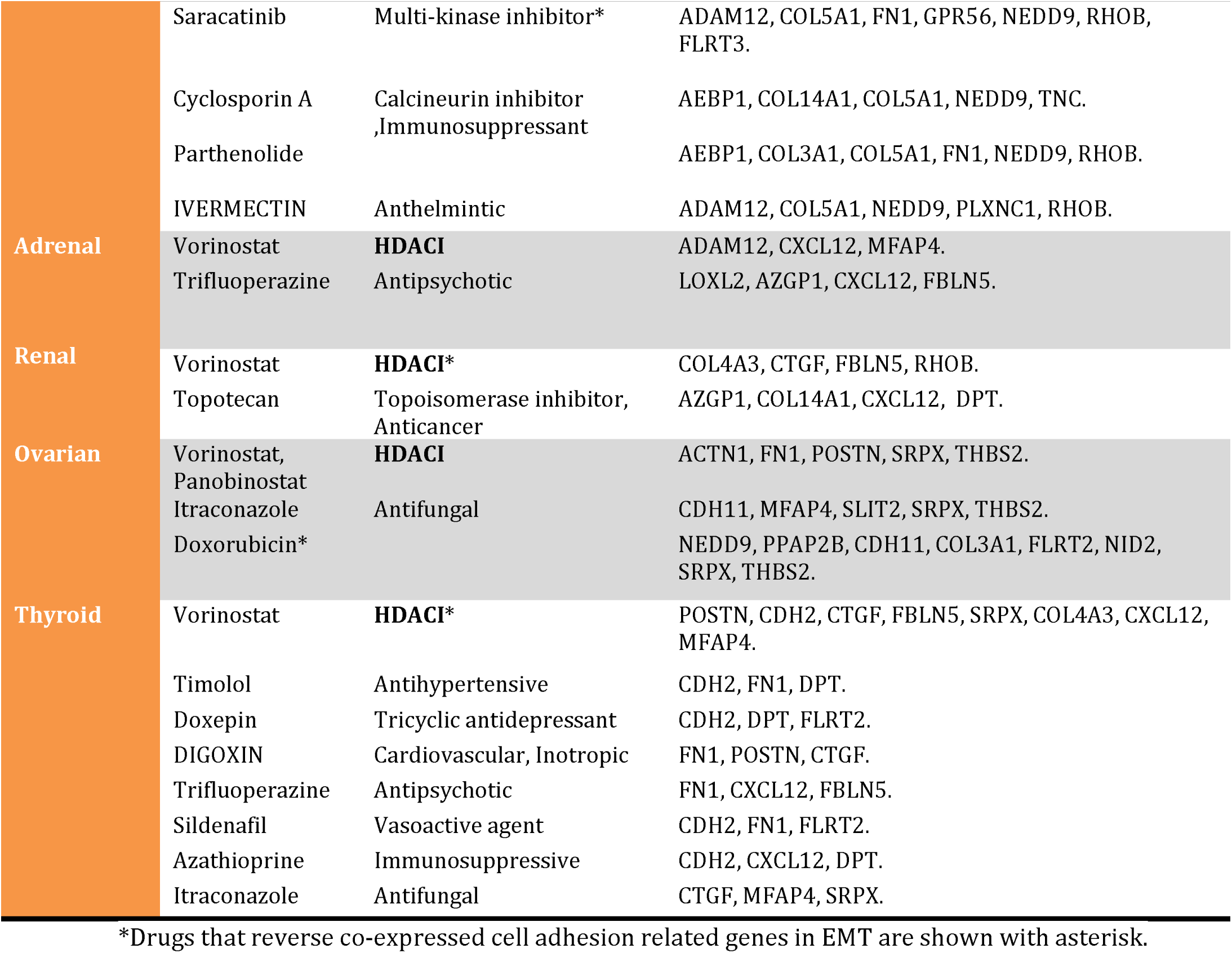
Drugs predicted by L1000CDS^2^ to reverse EMT gene sets in various cancers

To elucidate the underlying biology of the pan-cancer effects HDACIs on EMT, we hypothesized that these drugs may modify the epigenetic mechanisms governing mesenchymal plasticity related to stemness features and pluripotency which are conserved across cancers [42]. Cellular plasticity is a corollary of EMT-MET (Mesenchymal-Epithelial Transition) conversions in embryonic development with which early epithelial cells are reprogrammed to migrate during gastrulation, the critical step establishing the three germ layers and the body plan. Soundararajan et al. developed a 160-signature of embryonic plasticity signature from epiblasts at day 6.5 which represented the highest plasticity status mirroring the capacity of tumors for distant metastasis and patient survival [43]. The search for plasticity reversing drugs through L1000CDS2 returned Vorinostat to reverse expression of 13 genes related to plasticity (P-value in hypergeometric test =0.011) while this drug did not affect the expression of a664 adult tissue gene -signature which did not contain any of the embryonic gene signature **(Figure 3C)**. Accordingly, downregulated genes by Vorinostat in multiple cell lines were significantly enriched (FDR< 0.05) in stemness-related gene sets in MSigDB **(Figure 3D)**. Most significantly downregulated genes by Vorinostat were found to be significantly expressed in stem cells and stromal fibroblast cells compared to their expression profile in mouse hematopoietic cell strains deposited ImmGen database (Figure 3E). These results pinpoint the role of epigenetic plasticity in aggressive behaviors of carcinoma cells which would be inhibited by Vorinostat. Since a strong correlation between expression of embryonic plasticity gene signature and poor survival due to metastasis was observed in this study [43], we were also interested to see if the 13 plasticity-related gene set reversed by Vorinostat were associated with survival with DRUGSURV database which connects targets of drugs with survival. We observed that higher expression AnnexinA6 (ANXA6) was linked with increased survival in breast and high grade glioma cancer data sets which was increased by Vorinostat. Conversely, MYC, PTGIS, PUS7, SDC, and WDR12 expression were associated with decreased survival in various cancers and were downregulated Vorinostat **(Table 2)**.

**Table 2.** Association of plasticity-related genes with overall survival in DRUGSURV database that are reversed by Vorinostat

| Genes | Cancer type | Datasets | FDR adjusted P-value | Effect on Survival * |
| --- | --- | --- | --- | --- |
| **ANXA6** | Breast cancer | GSE25065 | 0.00025 | Positive |
|  | High-grade glioma | GSE4271 | 0.0023 |  |
| **MYC** | Breast cancer | GSE22226 | 0.0019 | Negative |
| **PTGIS** | Glioblastoma | GSE13041 | 0.00048 | Negative |
| **PUS7** | Lung cancer | GSE13213 | 0.00000973 | Negative |
|  | Breast cancer | GSE25055 | 0.0011 |  |
| **SDC1** | High-grade glioma | GSE4271 | 0.00018 | Negative |
| **WDR12** | Multiple myeloma | GSE24080 | 0.00038 | Negative |
|  | Chronic lymphocytic leukemia | GSE39671 | 0.0012 |  |
|  | Colon Cancer | GSE17536 | 0.0024 |  |
| **HDAC2** | Breast cancer | GSE22226 | 0.00255 | Negative |
|  |  | GSE21653 | 0.0204 |  |
|  |  | GSE31448 | 0.025 |  |
|  |  | GSE22220 | 0.0311 |  |

\*Positive and negative effects indicate increased and decreased patient survival following gene overexpression respectively.

### 3.3. Vorinostat treatment alone is not sufficient to inhibit dispersion in carcinoma cells

In order to experimentally validate the anti-EMT efficacy of hypothesized drugs, it is nontrivial to consider the transient nature of EMT as well as its dependency upon signaling from surrounding microenvironment and basement membrane [44]. We thus implemented a 3D microfluidic device with collagen embedded A549 lung carcinoma spheroids in co-culture with HUVEC endothelial cells to induce EMT and monitored dispersion of carcinoma cells from the spheroids for 84 hrs following drug treatment. This measurement was formerly shown to correlate with upregulation of Vimentin and downregulation of E-cadherin as hallmarks of EMT [40] and recapitulates the transient and dynamic nature of EMT following the real-time monitoring of drug effects in a physiologically relevant model. Treatment of A549 cells with 1µM Vorinostat did not show a significant decrease in cell dispersion from the collagen spheroids compared to control after 84 hrs **(Figure 4A, B)**. However, migration of HUVEC cells from the lateral channel of the devices toward A549 was less observed at the interface of disseminated carcinoma cells and the proliferated HUVECs **(Figure 4C)**. As we previously showed, Vorinostat affects cellular plasticity and given that this phenomenon in carcinoma cells is congruent with a permissive microenvironment, we analyzed the biological processes in A549 which were affected by Vorinostat. Vorinostat gene signature was enriched with processes involving cell migration, chemotaxis and angiogenesis. It could also decrease the expression of COL1A1, COL4A1, MMP2, TIMP2, LOXL1, CDKN1, HTRA1 and RAB31 which are associated with activated fibroblasts and stromal cells [45-47]. Accordingly, 24% of HDACI-induced gene products were located in extracellular vesicular exosomes and in extracellular matrix which affected cell adhesion, cell migration, chemotaxis, epithelial differentiation, angiogenesis and regulation of NF-kB kinase expression which are known to be associated with EMT **(Figure 4D,E)**.

**Fig.4.**
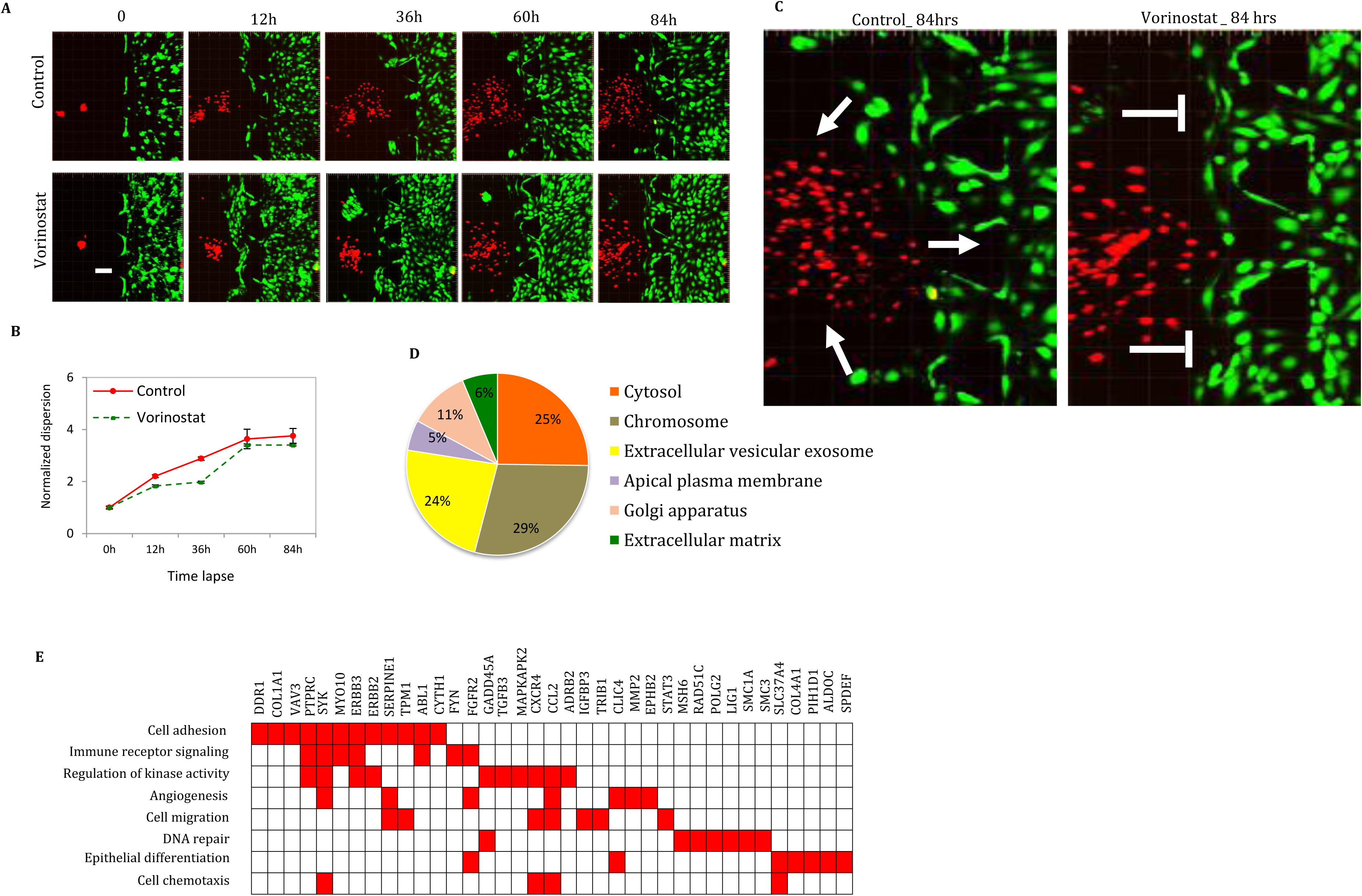
Vorinostat mono-therapy alone does not effectively in inhibit cell dispersion. **(A)** Confocal live cell microscopy images showing dispersion of A549 lung cancer cells (with red fluorescent) co-cultured with HUVECs (green fluorescent) following treatment with 1µM vorinostat versus control in microfluid device as a metric of EMT. Scale bar indicates 100 um. **(B)** Normalized time-lapse dispersion distance of A549 cells quantified by pixel distance measurement in MATLAB in Vorinostat or no treatment control group normalized by dividing dispersion distance at each time-point to 0h. Data are represented as mean ±SEM for three spheroids in two independent biological replicates. t-test was performed to assess statistical difference between Vorinostat and control at 84 hrs and P-value < 0.05 was considered significant. **(C)** Migration of HUVEC cell toward dispersed A549 cells in presence and absence of Vorinostat after 84 hours. **(D)** Cellular location of vorinostat-induced gene products using Enrichr. FDR adjusted P-value (q-value) <0.05 was considered significant for cellular component enrichment. (E) Significantly enriched (q-value < 0.05) biological processes for shared genes between collated EMT signatures and Vorinostat-induced differentially expressed genes (DEGs) from iLINCS database.

### 3.4. Elucidation of kinase-interaction network in EMT and their drug inhibitors

The inefficacy of Vorinostat to fully abrogate cell dispersion in the 3D environment clearly demonstrated the contribution of other intracellular pathways in EMT. As EMT is a stepwise process, we further analyzed the reversing drugs for time-course data delineating early and late differentially expressed genes by SNAIL upregulation [48]. The results returned Trichostatin-A, Dasatinib (SRC Kinase inhibitor) and Ketoprofen (a non-steroid anti-inflammatory drug) as first, fifth and six ranked drugs respectively to reverse early phase genes in EMT and Trichostatin-A as seventh ranked drug to reverse late phase genes in EMT. These results not only underscore the significance of overcoming epigenetic barriers to switch from epithelial traits into mesenchymal features independent of the cancer type [49, 50], but also underline the contribution of multiple cellular kinases in accomplishment of EMT. We thus sought to determine EMT-associated kinome by performing kinase-substrates enrichment analysis on the EMT library using “Expression2Kinases” software. The identified kinases were connected using “STRING 10.0” database to create a kinase interaction network (Supplementary File 3). The network was consisted of four highly connected sub-communities (modules) regulating different pathways in EMT (Supplementary File 4). Calculation of main network centralities proper for our network [51] including degree, betweenness and closeness showed that the proto-oncogene SRC with 45 interactions was the hub node (a node with high degree of interactions) in the whole network. SRC also owned the highest betweenness (0.27) as a measure of shortest paths which pass through this node and also highest closeness value (0.65) implying the role of SRC to be at the crossroad of multiple pathways (Figure 5A). Moreover, IKBKB owned the second rank for betweenness value (0.15) and the third rank for closeness (0.56) centrality in the network confirming its deliberate position in the network. IKBK was directly connected to SRC, TGF-beta, MAPK14 and CDK7 which were distributed among four other modules (Figure 5B). These results suggest that by dual targeting of SRC and IKBK, the whole kinase interaction network associated with EMT would be affected (Figure 5C).

**Fig.5.**
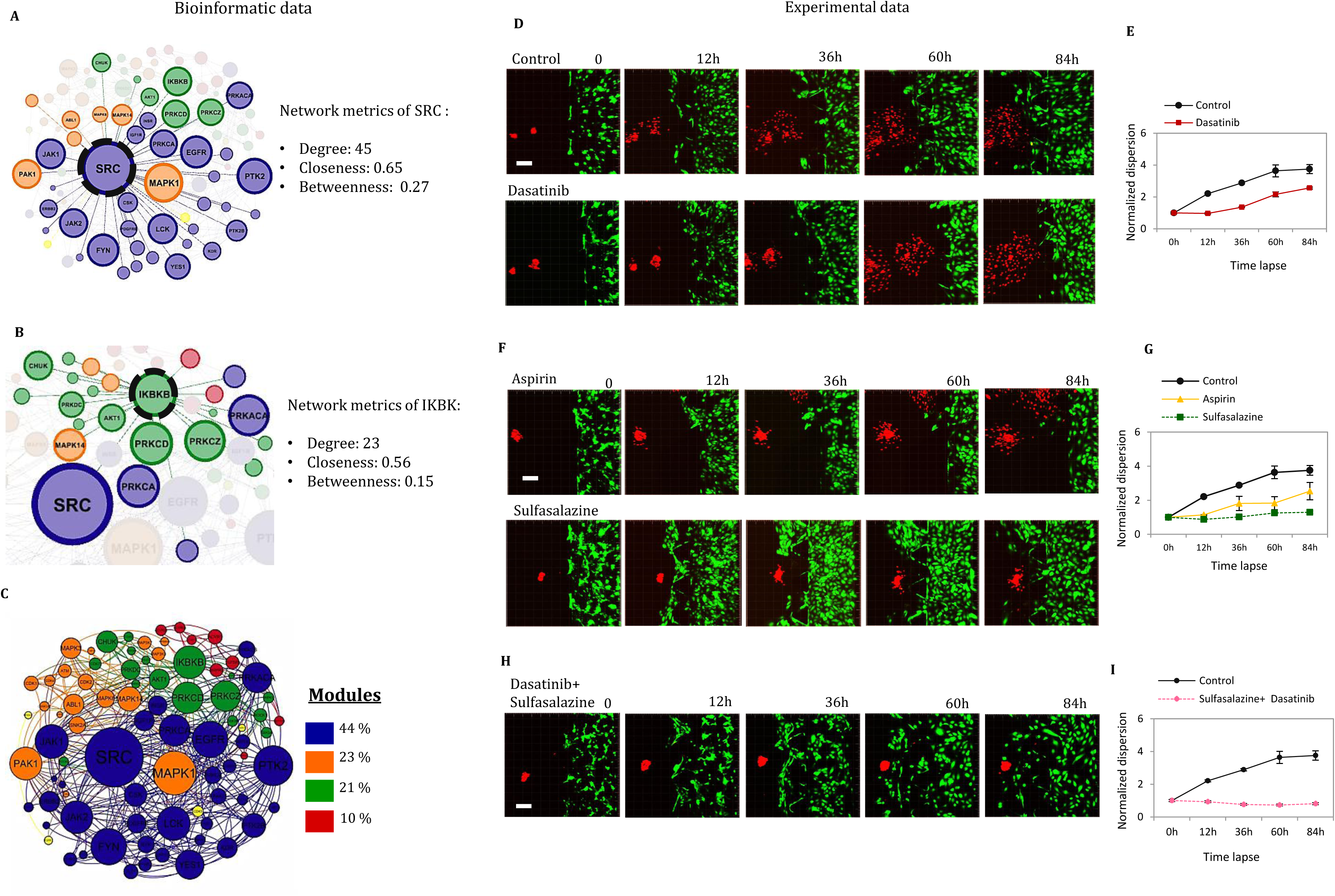
Kinase interaction network analysis identifies SRC and IKBK as principal druggable kinases in EMT to perturb the whole network. **(A)** Identification of SRC as a hub node and its first neighbor kinases in EMT-associated kinase interaction network based on centrality metrics degree, betweenness and closeness. **(B)** Identification of IKBK as another principal kinase in EMT functionally connected to 3 sub-communities in the network. **(C)** Overall view of Kinase-Interaction Network with its constituent modules that is perturbed by co-targeting of SRC and IKBK. Distinct modules are given separate colors **(D)** Experimental results of A549 dispersion in spheroids after treatment with 1 µM Dasatinib as a SRC kinase inhibitor for 84 hrs co-cultured with HUVECs. **(E)** Measurement of dispersion distance of A549 cells from collagen spheroids in Dasatinib and no treatment control normalized by dividing to distance at 0 time point. **(F)** Treatment of A549 spheroids and monitoring cell dispersion with 1 µM Aspirin or Sulfasalazine for 84hrs as IKBK inhibitors. **(G)** Normalized dispersion distance following treatment with Aspirin or Sulfasalazine for 84 hrs. **(H, I)** Significant (t-test, P-value< 0.05) inhibition of A549 dispersion distance normalized to 0h time-point from the spheroids following co-treatment with Dasatinib and Sulfasalazine at 84hr. Data represent mean ±SEM for three spheroids in two biological replicates. Scale bar indicates 100 um.

### 3.5. SRC and IKBK overexpression leads to induction of EMT in epithelial cells

To confirm the relevance of SRC and IKBK kinases in EMT, ***“CREEDS”*** database of gene perturbation signatures [37] was queried and differentially expressed genes (DEGs) by SRC and IKBK overexpression or constitutive activation were identified. Biological functions of DEGs in the SRC overexpression signatures were associated with cell adhesion and actin cytoskeleton organization. In IKBKB over-activation signatures, epithelial cell differentiation, extracellular matrix organization, cell junction organization, regulation of inflammatory response were observed (adjusted P-value<0.05), all of which are evidently associated processes in EMT. Consistently, the overlap between DEGs in the SRC and IKBK perturbation signatures and the 962-EMT gene set was significant (P-value< 0.01) in hypergeometric statistical test **(Supplementary File 5)**.

### 3.6. Combination of SRC and IKBK inhibitors with Vorinostat reduces carcinoma cell dispersion and endothelial cell invasion

To identify Food and Drug Administered (FDA)-approved drugs that inhibit SRC and IKBK targets, EMT-associated kinases were mapped onto Drug-Target Network [52, 53]. Four approved drugs including Dasatinib, Bosutinib, Nintedanib and Ponatinib were found to inhibit SRC kinase family along with multiple other kinases such as PDGFR. Aspirin, Sulfasalazine, Mesalazine currently used for treatment of inflammatory diseases were also found to inhibit IKBK. We thus chose Dasatinib with 23 targets including SRC kinase family and Aspirin with 11 targets including IKBK for experimental confirmation of their anti-EMT effects. Inhibition A549 cells dissociation from the spheroids was not statistically significant following treatment with 1µM Dasatinib alone for 84 hrs compared to control no-treatment group **(Figure 5D,E)**. Independently, Aspirin alone at 1µM concentration did not significantly inhibit dispersion. Moreover, 1µM Sulfasalazine, another identified IKBK inhibitor which also inhibits CHUK kinase, imposed a delay on A549 dispersion for about 72 hrs compared to negative control **(Figure 5F,G)**. Based on the insights from kinase interaction network analysis however, combination of 1µM Sulfasalazine with Dasatinib significantly inhibited cell dispersion from the spheroids in the co-culture **(Figure 5H,I)**. Interestingly, addition of Vorinostat to combination of Dasatinib and Aspirin, with a low concentration of 1µM of each drug, fully inhibited cell dispersion **(Figure 6A, B)** and a distance of 200 µm (equal to the space between the inputs of the two channels of the device) was kept between the two co-cultured cell types **(Figure 6C)**. It was also grossly evident from green pixel count in the images that the triple-combination treatment maintained survival of co-cultured HUVEC endothelial cells which denotes tolerability of proposed combination regimen for normal cells **(Supplementary File 6)**

**Fig. 6.**
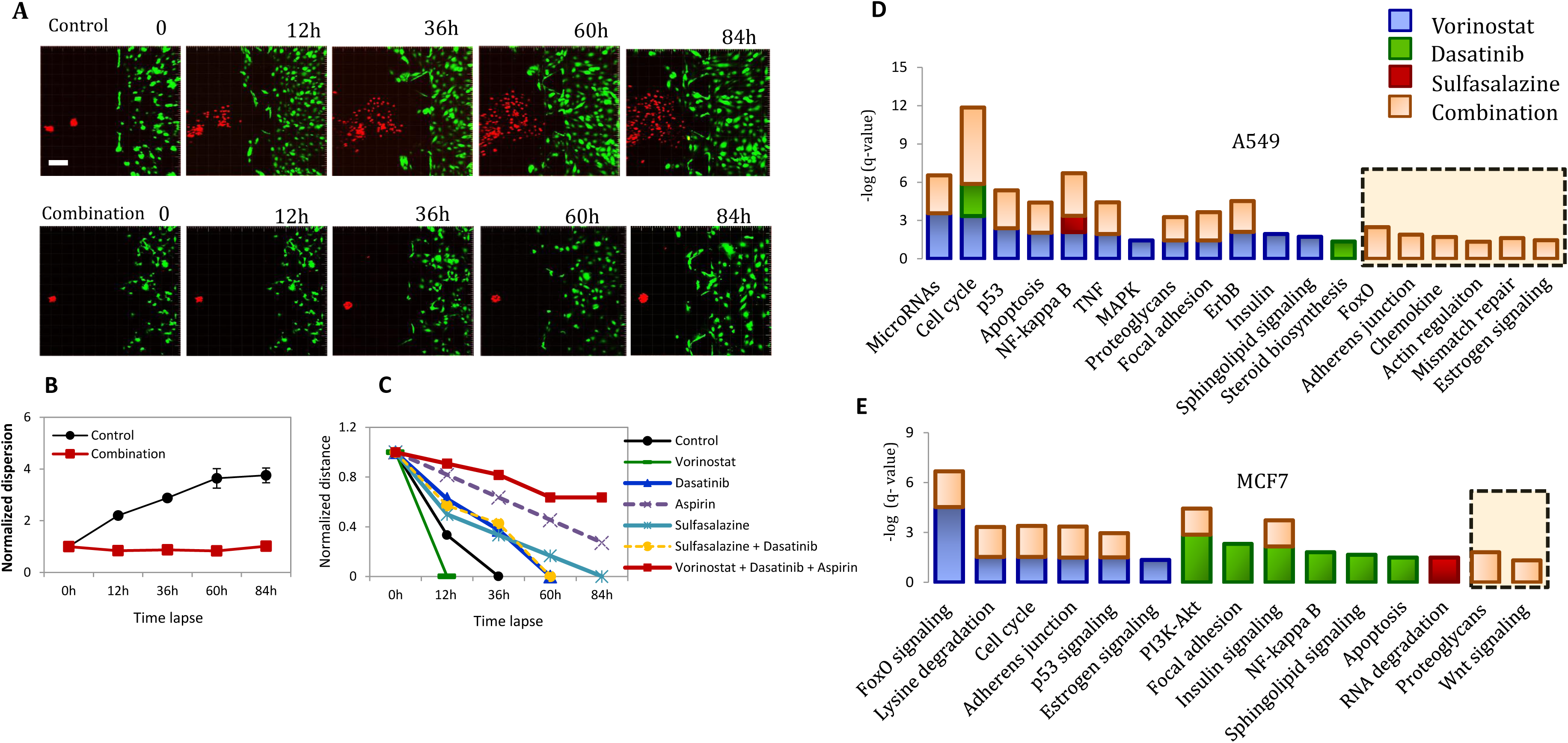
Combinatorial treatment with Vorinostat, Dasatinib and Aspirin inhibits A549 cell dispersion and invasion by affecting multiple pathways. **(A)** Assessment of cell dispersion following combination treatment with 1 µM of each drug for 84 hrs. Scale bar length represents 100 um **(B)** Significant (t-test, P-value < 0.05) dispersion distance between null control versus combinatorial triple treatment at 84hr was observed. Data represent mean± SEM and the error bars indicate dispersion measurements in two independent spheroids. **(C)** Distance between dissociated A549 and HUVEC cells normalized to 0hr as a measure of endothelial cell migration toward carcinoma cells **(D)** KEGG Pathway enrichment analysis in A549 and **(E)** MCF7 cells for differentially expressed genes following treatment by HDACIs, kinase inhibitors and their combination obtained from iLINCS database. Significantly enriched pathways are determined by FDR-adjusted q-value < 0.05). Unique pathways observed with drug combination are marked with dashed value. Color codes represent each drug and the combination.

### 3.7. Analysis of EMT alterations by combination of HDACIs and kinase inhibitors

To elucidate the underlying pathways leading to significant inhibition of EMT by triple drug combination, gene expression changes of Vorinostat, Dasatinib and Sulfasalazine were extracted from ***“iLINCS”*** for A549 and MCF-7 carcinoma cell lines. Expression changes for three drugs were also combined in each cell line to estimate the affected pathways following drug combination. Several pathways involved in EMT were enriched by combination of DEGs. In A549, a lung adenocarcinoma cell line with reversible mesenchymal traits, the combinatorial drug signature was associated with manipulation of several EMT-related pathways including adherens junction and regulation of actin cytoskeleton, Foxo signaling, chemokine and NF-kB signaling **(Figure 6D)**. Other pathways including mismatch repair and cell cycle were also affected by drug combinations which are underlying mechanisms of EMT-induced drug resistance. However, in MCF-7, a hormone responsive breast carcinoma cell line with epithelial traits only cell cycle, proteoglycans and Wnt signaling pathways were enriched in the combination signature **(Figure 6E)**. To further confirm the relevance of identified drug targets in EMT and aggressiveness of cancers, expression of drug combination targets were analyzed in ovarian cell lines in the NCI60 panel. An EMT score was attributed to each cell line by subtracting the z-score expression of Vimentin from CDH1 as these genes showed the most anti-correlated mesenchymal and epithelial markers in NCI60 panel respectively (r= -0.7) so that cell lines with positive score were assumed as mesenchymal. The z-score expressions of drug targets with relevance to EMT were then extracted from CellMiner and were compared across mesenchymal ovarian cell lines. A positive correlation (r >0.5) between increase in the EMT score and expression of FYN (a member of SRC family of kinases), PDGFRB, HDAC2, CHUK and IKBK which are all inhibited by our proposed drug combination was observed implying that by increasing mesenchymal properties in these cell lines, the expression of these drug targets increased. To see if the pattern was maintained in datasets of clinical samples, the association of these drug targets with various histopathological characteristics of ovarian cancers in CSIOVDB database was assessed **(Supplementary file 7)**. Among the drug targets, FYN and PDGFRB, CHUK and HDAC2 were correlated with mesenchymal subtype. Moreover, expression of PDGFRB was associated with transition from stage 3 to 4 and was more enriched in mesenchymal, stem-A and Stem-B subtypes of breast cancer. Higher expression of CHUK and PDGFRB were associated with decreased survival. From the indirect drug targets with decreased expression in stromal cells by Vorinostat, expression of COL1A1, MMP2, TIMP2 and HTRA1 were enriched in mesenchymal subtype of ovarian cancer among them, expression of LOXL1 and MMP2 were also associated with decreased survival. These results indicate that the triple drug combination that we proposed inhibits multiple aspects of EMT, tumor invasion and could ultimately improve patient survival including regulation of 1) extracellular matrix remodeling 2) cell adhesion and motility, 3) cell cycle related pathways and 4) inflammatory pathways in mesenchymal-like cells and rewiring stromal cell reprogramming and 5) inhibition of tumor cell plasticity and stemness.

## 4. Discussion

Here we practiced a systems pharmacology methodology by integrating currently available bioinformatic resources with network analysis to translate genome-wide EMT expression signatures into a predictive pipeline of drug combination design. Despite the emergence of targeted therapies for cancer, long-term regression of the disease is hindered by development of resistance emanating from feedback regulatory mechanisms and redundancies embedded in signaling pathways with cross-talks. Hitting such robustness requires a rational combinatorial design rather than naïvely screened combinations for complex phenotypes such as EMT in carcinoma [54].

Following analysis of cell-adhesion related genes in EMT for each carcinoma type, we observed HDACIs as a consistent drug class to down-regulate expression of EMT-associated genes in carcinoma types analyzed in this the study. Confirming these results, Tang et al. observed that among multiple drug classes, HDAC inhibitors could restore E-cadherin expression with low cytotoxicity [55]. We also showed that HDACIs can potentially affect EMT through inhibiting the plasticity and communication of tumor cells with their microenvironment. From the clinical point of view however, mono-therapy with HDACIs did not completely control the disease state at metastatic stage however their combination with chemotherapy regimens or targeted drugs seemed to be significantly effective in disease remission [56]. Such discrepancy at the clinical level was reflected in our computational as well as experimental sections of our study in 3D co-culture system since as we showed, HDACIs inhibit only part of the many mechanisms leading to EMT and their efficacy was potentiated when added to intracellular kinase inhibitors [57].

To find other appropriate co-targets along with epigenetic modulating drugs, we used network biology to rank the principal kinases in the execution of EMT. Many of the identified kinases in the constructed network have already been proposed as druggable targets to inhibit EMT in separate parts of literature including EGFR, PGFRR, EPHB2, YES1, LCK, and AXL [58-62]. While it is not possible to inhibit all such redundant kinases in clinical settings, mathematical analysis of kinase interaction network showed the addressed kinases are functionally connected to SRC family of kinases which owned the highest degree and betweenness values in the network making them appealing druggable targets on which the information flow from the addressed upstream kinases is converged on. Consistently, Avizienyte et al. showed that accumulation of SRC kinase is associated with EMT [63] and SRC kinase inhibitors were able to inhibit metastasis to liver, breast and colon cancer cells by restoring the expression of E-cadherin [64]. However, search in the literature shows that despite the initial excitements in potentiality of SRC kinase inhibitors to inhibit metastasis, the results were not promising in later clinical stages [65]. We similarly found that in 3D co-culture environment, low dose treatments with Dasatinib as a SRC kinase inhibitor alone did not inhibit cell dispersion from the initial aggregates.

Closer inspection of the kinase-interaction network revealed that by targeting SRC kinase alone, only three modules of the network (shown with blue, orange and green nodes) were manipulated while an important subnetwork associated with cytokine-cytokine receptor interaction and TGF-beta signaling (red nodes) was unaffected. The kinase interaction network elucidated that IKBK interacts with ROCK1, AKT1, GSK and CHUK (IKK-α) kinases which are dispersedly mentioned in the literature to fulfill EMT [66-69]. Consistent with these results, we observed EMT inhibition when IKBK was co-targeted beside SRC and their combination with HDAC inhibitors was effective in limiting EMT consequences. Addition of Vorinostat to this combination also confined invasion of endothelial cell towards spheroids and maintained the inhibition of dispersion from the spheroids. In line with this observation, Gopal et al. showed that aggressive cancer cells undergoing EMT induce angiogenesis by secreting vesicles which we showed to be inhibited by Vorinostat [70].

In the end, by implementing a 3D co-culture system to simulate tumor microenvironment and use of clinically achievable doses of already prescribed drugs, we observed agreement between in silico and in vitro results to confirm the relevance of proposed combinatorial regimen in abrogating a process such as EMT. Such modeling assisted in transfiguring segmented data in the literature into a hypothesis-generation tool for another round of in vivo and clinical research. Moreover, given the safety of the proposed drugs in this study, as well as their FDA approval for other indications, the outcomes presented here, are of direct translatability to clinical trials.

## Acknowledgement

Contributors to publicly available open source tools are greatly appreciated. We also acknowledge Prof. Avi Ma’ayan for helpful comments. This manuscript was not performed with public or institutional grants.

## Competing interests

Authors declare no confliction in the interest.

## Supplementary Files

Supplementary File 1. Meta-analysis results of up and downregulated genes across various cancers which are related to cell adhesion (GO:0007155) which were performed in CancerMA database.

Supplementary File 2. Expanding upregulated and downregulated genes in meta-analysis with patients data in TCGA using co-expression module in cBioportal. Pearson’s coefficient >0.8 was considered significant.

Supplementary File 3. The edge list for creating kinase interaction network.

Supplementary File 4. Common genes between genes that are differentially expressed in SRC and IKBK overexpression (OE) or overactivation (OA) and are observed in EMT signatures as well.

Supplementary File 5. KEGG pathway enrichment analysis for kinases within each four module of kinase interaction network. Hub kinases which are nodes with highest degree within each module are also showed.

Supplementary File 6. Green pixel count with MATLAB representing number of HUVEC cells in time-points between 0 to 84 hrs following treatment with proposed drugs and their combinations.

Supplementary File 7. A) The correlation between EMT score and expression of drug targets in ovarian cells in NCI-60 panel of cancer cell lines. B) Among the drug targets in patient samples, PDGFRB expression is positively correlated with EMT score. C) The association between direct targets of proposed drugs with EMT score and patient survival. D) Expression of the targets in subtypes of ovarian cancer in CSIOVDB” database of ovarian cancer microarray gene expression version 1.0.

